# In Silico Post-screening of Anti-polymerization Agents to Treat Sickle Cell Disease

**DOI:** 10.1101/2025.10.25.684554

**Authors:** Ying Qian, Nazanin Ahmadi Daryakenari, Melissa Hallow, Lu Lu, Pierre A. Buffet, Ming Dao, He Li, George Em Karniadakis

## Abstract

Sickle cell disease (SCD) is a genetic disorder that affects approximately 100,000 individuals in the United States and millions globally. Although curative therapies, such as stem cell transplant and gene therapy, are available, their application is limited by high cost and donor availability. Consequently, drug therapy remains the most feasible treatment option for the majority of patients with SCD. To date, only four drugs have been approved by the FDA, but none of these treatments comprehensively address all SCD-related symptoms or crises, suggesting the pressing need for developing new drugs. Several high-throughput screening campaigns have been performed for SCD drug discovery based on the *in vitro* sickling of red blood cells (RBCs) and they have identified several hits. However, it is challenging to replicate the organ-specific oxygen level and physiological deoxygenation time in these *in vitro* RBC sickling assays. Furthermore, these assays do not consider the pharmacokinetics (PK) and pharmacodynamics (PD) of the identified drugs, which are essential to determine whether the drugs can provide robust and sustained efficacy in treated patients with SCD. To address these technical gaps, we have developed a computational platform to perform post-screening analysis of potential anti-sickling agents. This platform sequentially combines PK/PD models with a kinetic model of RBC sickling, enabling efficient prediction of the dosage-dependent therapeutic efficacy of various anti-sickling agents based on patient-specific hematological factors and organ-specific oxygen levels. We first demonstrate the effectiveness of our integrated platform by showcasing the therapeutic efficacy of two FDA-approved drugs, Hydroxyurea (HU) and voxelotor. Next, we evaluate the therapeutic efficacy of two potential anti-sickling agents under clinical trial, namely Bitopertin and osivelotor. Our findings suggest that Bitopertin exhibits anti-sickling effects that are considerably less pronounced than those of HU and voxelotor. On the other hand, osivelotor can achieve similar anti-sickling effects as voxelotor with significantly lower doses due to its improved PK properties. Furthermore, we show the versatility of the proposed platform in predicting the anti-sickling effect of multi-agent therapies and evaluate the consequences of drug noncompliance. In particular, our analysis indicates that noncompliance with voxelotor may result in rapid increases in RBC sickling, whereas osivelotor is likely to mitigate noncompliance-induced adverse effects due to improved PK properties. We further quantify the relationship between drug dosage and the duration of noncompliance that leads to loss of therapeutic efficacy for voxelotor and osivelotor, providing guidance for optimizing dosage strategies to reduce the risk associated with noncompliance. In summary, our *in silico* platform serves as a valuable tool for post-screening analysis of potential anti-sickling agents by considering their PK and anti-sickling efficacy under patient-specific hemoglobin level and organ-specific oxygen level, thereby gaining insights into their potential therapeutic efficacy alone or in combination before clinical trials.

## 1 Introduction

Sickle cell disease (SCD) remains a significant global health burden, particularly in sub-Saharan Africa, India, and the Middle East, where limited healthcare access contributes to high childhood mortality. An estimated 300,000 infants are born with SCD globally each year [1]. SCD originates from a single-nucleotide mutation in the hemoglobin gene, leading to the synthesis of hemoglobin S (HbS) instead of normal hemoglobin A (HbA) during erythropoiesis. Hemoglobin, the oxygen-carrying protein within red blood cells (RBCs), undergoes a conformational change from the relaxed (R) state to the tense (T) state upon deoxygenation that predisposes HbS to polymerize into HbS fibers [2–4]. As these fibers expand, they distort RBCs and impair their flexibility, ultimately leading to vaso-occlusive crises (VOCs), a hallmark of SCD. Recurrent and unpredictable vaso-occlusion can precipitate severe complications, such as acute chest syndrome (ACS), splenic sequestration, and nephropathy [5, 6]. In addition, although sickled cells can revert to a normal shape when reoxygenated, the repeated cycles of sickling and unsickling hasten hemolysis, shortening the lifespan of RBCs to an average of 64.1 days (range 35–91 days) [7, 8] in comparison to *∼*120 days for healthy RBCs, thereby contributing to anemia, a common clinical manifestation of SCD.

Advancements in understanding SCD pathophysiology have led to the development of various treatments [9]. Current treatment strategies for SCD include stem cell transplant [10–13], gene therapy [14–16], transfusion and medications [17–19]. Stem cell transplants and gene therapies are curative, but their implementation will not be universal due to either the availability of the donors, risks associated with intensive myeloablation, or the high complexity of manufacturing. Blood transfusion requires a consistent and accessible blood supply while posing risks such as iron overload when other methods than exchange transfusion are used. Thus, drug therapy remains a more feasible choice for most SCD patients, particularly in economically disadvantaged countries. Four drugs, namely hydroxyurea (HU), voxelotor, crizanlizumab, and L-glutamine, have been approved by the FDA for treating SCD. While L-glutamine and crizanlizumab alleviate the downstream sequelae of SCD, they do not improve hematologic parameters, i.e., anemia. On the other hand, HU, targeting the root cause of RBC sickling, is beneficial in SCD by reducing anemia and the frequency of VOCs, and ACS [20, 21]. Nevertheless, its efficacy in preventing stroke, nephropathy, and pulmonary hypertension is not evident. Voxelotor, another anti-sickling agent, mitigates anemia [22, 23], but its impact on the rates of VOCs or ACS was not demonstrated. Not least, voxeletor was withdrawn from the market in 2024 due to safety concerns, possibly related, in some cases, to the stability of its effect in poorly compliant patients. There is therefore a pressing need to develop new drugs.

Drug development is a time-consuming and multi-stage process. A more efficient strategy for identifying SCD therapies focuses on repurposing compounds that are already FDA-approved for other diseases or have previously undergone clinical trials, given their demonstrated anti-sickling activities. This approach allows these agents to move into clinical trials without the prolonged preclinical studies and early clinical studies that are mandatory to ensure efficacy and safety before wide administration to human subjects. Several high-throughput sickling-based screening assays have been built for SCD drug discovery [24–26]. In particular, Eaton’s team at the NIH established a high-throughput sickling kinetics assay to screen the Scripps ReFRAME drug repurposing library and identified hundreds of potential SCD drug candidates [27]. However, these *in vitro* assays do not fully account for the pharmacokinetics/pharmacodynamics (PK/PD) of the drugs, and the experimental conditions involve an extreme 0% oxygen level to induce RBC sickling, which does not accurately reflect the physiological oxygen environment in human tissues. On the other hand, *In silico* modeling [28–40] has been used as a complementary tool in unraveling the pathogenesis of SCD and filling the knowledge gaps of traditional *in vitro* and *in vivo* approaches. These computational models could be particularly valuable for studying diseases like SCD, where patient-specific factors and physiological conditions, such as oxygen gradients, significantly influence drug response [41, 42]. Finally, because co-administration of multiple drugs will likely enhance efficacy, modeling the impact of multi-drug regimens should again notably accelerate the identification of the most promising drug combinations to reduce end-organ complications in SCD.

In this study, we propose the development of a computational platform designed to perform post-screening analysis, enabling the down-selection of potential anti-sickling agents identified through high-throughput screening. This platform sequentially pipelines the PK/PD analysis and a sickling kinetic model that allows predictions of RBC sickling by accounting for the patient-specific hematological parameters and organ-specific oxygen levels. The embedded kinetics model explicitly describes the kinetics of both homogeneous and heterogeneous nucleation of HbS monomers using nucleation theory while employing chemical rate laws to model HbS fiber growth dynamics [32]. Unlike prior frameworks that independently evaluate either PK/PD [18] or sickling kinetics [32, 43, 44], our platform uniquely integrates both, along with tissue-specific oxygenation, to enable mechanistic, patient-specific predictions of drug efficacy under realistic dosing and physiological conditions. Our comprehensive pipeline quantitatively assesses the feasibility of potential anti-sickling agents and evaluates their efficacy across various dosage regimens over time. We first validate the proposed computational platform by confirming the PK/PD mechanisms associated with sickling reduction of two FDA-approved medications, namely HU and voxelotor. Subsequently, we leverage the system to investigate Bitopertin, a glycine transporter 1 inhibitor drug candidate currently under clinical evaluation for erythropoietic protoporphyria that exhibits anti-sickling effects for SCD treatment, such as decreases in mean corpuscular hemoglobin concentration (MCHC). Moreover, we demonstrate that the pipeline enables the quantitative evaluation of the therapeutic impact of administering two agents concurrently, as well as assessment of the consequence of medication noncompliance, which is critical to understanding the association between noncompliance and adverse clinical outcomes. This computational platform could deliver critical insights into the therapeutic potential of candidates prior to clinical trials, potentially accelerating the drug development process.

## 2 Data and Methods

This section outlines the data sources and computational models used to develop and validate our *in silico* platform for evaluating anti-sickling drug efficacy. We first describe PK/PD data used for model calibration, followed by an overview of the integrated computational framework.

### 2.1 Data

We curated PK/PD data for three anti-sickling drug candidates – Bitopertin, voxelotor and osivelotor – from published clinical and preclinical studies. These datasets were used to calibrate and validate the drug-specific components of the platform under both single- and multiple-dosing regimens.

#### 2.1.1 Bitopertin PK/PD Data

To calibrate the PK model for Bitopertin, we used plasma drug concentration data measured from normal subjects for both single and multiple dosages. The single-dosage data, sourced from [45], include plasma concentrations following oral doses of 3 mg, 12 mg, and 24 mg. In addition, multiple-dosage data obtained from [46] correspond to a 30 mg oral regimen. For PD model calibration, we adopted data from [47], consisting of MCHC levels measured over 240 days in a Phase-I clinical trial on normal subjects. During the first 120 days, normal subjects were divided into groups and administered multiple doses of 10 mg, 30 mg, or 60 mg. After treatment discontinuation at day 120, MCHC levels continued to be monitored.

#### 2.1.2 Voxelotor PK/PD Data

Voxelotor PK model calibration was based on RBC drug concentration data from both single and multiple dosage regimens measured from normal subjects and SCD patients, sourced from [48]. The single-dosage data include concentrations following oral doses of 100 mg, 1000 mg, 2000 mg, and 2800 mg. Meanwhile, multiple-dose data include RBC drug levels after oral administration of 300 mg, 600 mg, and 900 mg. For the PD model, we used *O*_2_ saturation data at several percentages of hemoglobin modification, as reported in [49].

#### 2.1.3 Osivelotor PK/PD Data

Osivelotor PK model calibration was performed using RBC drug concentration data from both single- and multiple-dose regimens in SCD patients, as reported in [50]. The single-dose data comprised RBC drug concentrations following a 100 mg oral administration, while the multiple-dose data included RBC drug concentrations for oral dosing of 50 mg daily for five weeks, immediately followed by 100 mg daily for three weeks. For the PD model, we utilized the percentage of hemoglobin occupancy at various dose levels reported in the same study.

### 2.2 Computational Models

The structure of the proposed computational pipeline for evaluating anti-sickling drug candidates is illustrated in Fig. 1. This comprehensive pipeline sequentially integrates PK/PD modeling, a kinetic model for simulating the HbS nucleation and polymerization, and a biomechanics model to determine the sickling of the RBCs. The primary objective is to simulate the dynamic effects of the drug over time. Through PK/PD modeling, we capture the drug’s absorption, distribution, metabolism, and its pharmacological impact on biological variables that affect the HbS polymerization and RBC sickling. The output from the PK/PD model is subsequently fed into the kinetic model, which is informed by a biomechanics model to predict the rate and extent of RBC sickling. This integrated approach enables a mechanistic understanding of how anti-sickling drug candidates influence disease progression and provides quantitative predictions to evaluate the efficacy of potential anti-sickling drugs.

**Figure 1.**
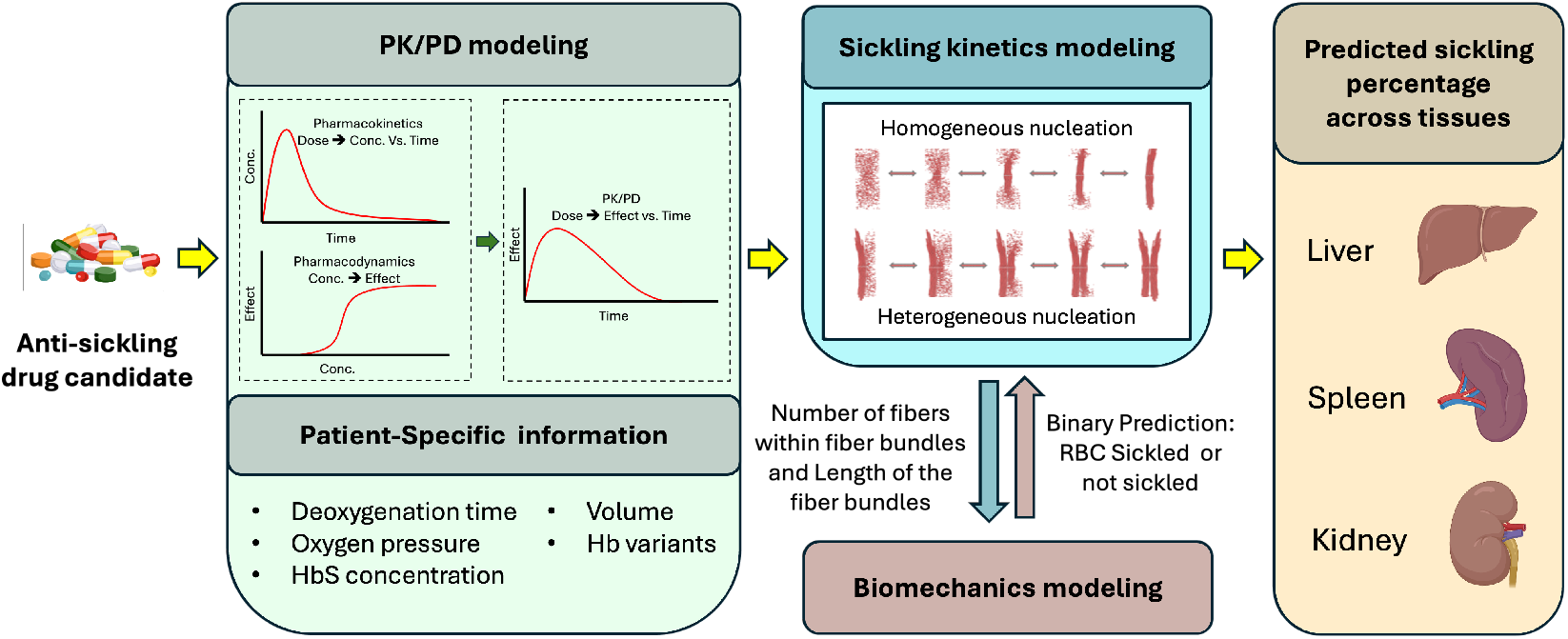
Overview of the proposed computational platform for evaluating anti-sickling drug efficacy. Starting from PK/PD modeling, the framework sequentially feeds into a sickling kinetics module and a biomechanics-based decision layer to predict tissue-specific RBC sickling fractions. The model incorporates patient-specific hematologic variables and organ-level oxygenation to enable personalized prediction of therapeutic response.

#### 2.2.1 PK Model

Since our platform’s main goal is to analyze the potential efficacy of anti-sickling agents rather than examine inter-patient or organ-specific variability in drug concentration, we have opted for basic PK/PD models instead of more advanced population PK/PD or physiologically based PK/PD approaches. Here, we use the same structure of the PK model for all drugs: Bitopertin, voxelotor, and osivelotor. Specifically, we selected a two-compartment PK model with oral dosing to characterize the drug’s absorption, distribution, and elimination. As described in Fig. 2A, this model consists of three components: a depot compartment representing the site of drug administration, a central compartment corresponding to systemic circulation, and a peripheral compartment reflecting drug distribution into tissues. The dynamics of this model are governed by the following system of ordinary differential equations (ODEs):

**Figure 2.**
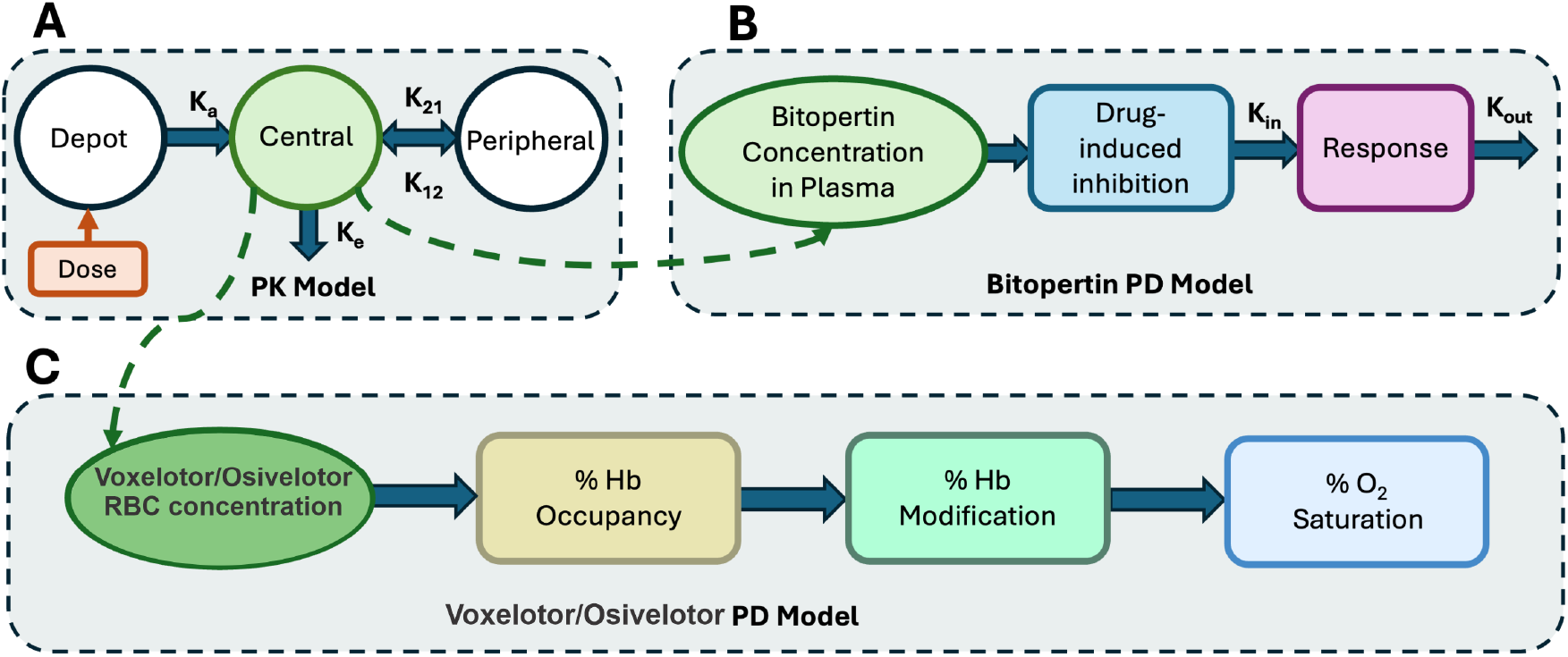
Schematic representation of the PK/PD modeling analysis that is connected with the sickling kinetics model to predict the sickling fraction of RBCs under drug effects. (A) A two-compartment PK model is used for Bitopertin, voxelotor and osivelotor. (B) For Bitopertin, an indirect response PD model captures its inhibitory effect on the production of the target response using the term 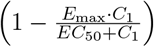. (C) The PD model for voxelotor and osivelotor involves a series of transformations: RBC drug concentrations from the PK model are converted into hemoglobin occupancy, then hemoglobin modification, and finally, the predicted percentage of oxygen saturation (*O*_2_ saturation).

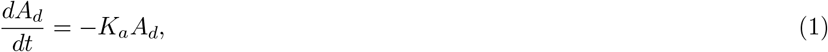

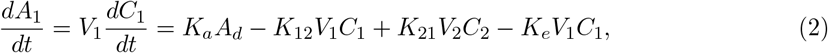

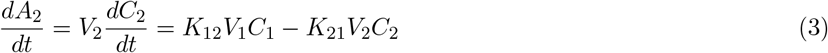

Given the oral route of administration, we model drug absorption using a two-compartment framework with a first-order absorption process. The depot compartment represents the gastrointestinal (GI) tract, from which the drug enters the central compartment (systemic circulation). Here, *A* denotes the amount of drug (mass), *C* the drug concentration (mass/volume), and *V*_*i*_ the volume of distribution in the *i*^th^ compartment (volume). The model includes the following first-order rate constants (units: 1/time): *K*_*a*_ for absorption from the depot to the central compartment, *K*_12_ and *K*_21_ for distribution between the central and peripheral compartments, and *K*_*e*_ for elimination from the central compartment. Specifically, *A*_*d*_, *A*_1_, and *A*_2_ represent the amount of drug in the depot, central, and peripheral compartments, respectively; *C*_1_ and *C*_2_ denote the corresponding concentrations; and *V*_1_ and *V*_2_ are the volumes of distribution in the central and peripheral compartments.

The initial conditions for the ODE system are given below, where *Dosage* denotes the amount of drug administered.

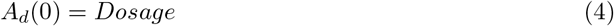

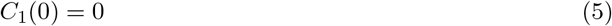

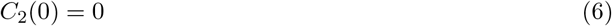

Overall, there are six parameters to be estimated in this ODE system, including *V*_1_, *V*_2_, *K*_*a*_, *K*_12_, *K*_21_, and *K*_*e*_.

Model fitting and parameter estimation for the PK model were performed using the `optim` function in R, which implements gradient-based optimization algorithms. The resulting system of ordinary differential equations was then solved using the `ode` function from the `deSolve` package to simulate the drug concentration-time profiles.

To quantify the uncertainty of the simulation results, we assumed the parameters follow a Gaussian distribution. We then generated 10,000 groups of parameter samples, using the estimated parameter values as the mean and the variances derived from the Hessian matrix returned by the `optim` function as the standard deviations. For each group of parameter samples, we solved the ODE system. Finally, we calculated the 5th, 50th, and 95th percentiles among the results at each time point.

#### 2.2.2 PD Model for Bitopertin

We aim to model how Bitopertin’s effect on MCHC levels changes over time through PD analysis. MCHC is a hematological parameter that reflects the concentration of hemoglobin per RBC, and it is highly associated with the intracellular hemoglobin polymerization and RBC sickling [51–53]. Given our prior knowledge that Bitopertin inhibits hemoglobin production, we chose to employ an indirect response PD model to capture this inhibitory effect on the production of the target response, as described in Fig. 2B. This model is particularly suitable for describing delayed drug effects mediated through intermediate physiological processes. The dynamics of this system are governed by the following ODE:

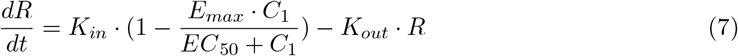

where 0 *< E*_*max*_ *<* 1. *E*_*max*_ denotes the maximum effect of the drug as the concentration goes to infinity with unit 1. *EC*_50_ denotes the plasma drug concentration that produces 50% of maximum inhibition with unit mass/volume. *C*_1_ denotes the plasma drug concentration with unit mass/volume. *K*_*in*_ denotes the rate constant for the production of response with unit 1/time. *K*_*out*_ denotes the rate constant for dissipation of response with unit 1/time.

The initial condition of the ODE is given below, where *R*_0_ represents the target response level under normal conditions.

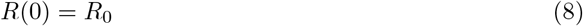

In the absence of any drug, the response is 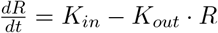. At steady state 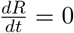, so it must be true that

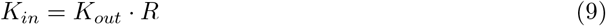

The process for solving the ODE system and quantifying the uncertainty of the simulations follows the same approach described in Section. 2.2.1.

#### 2.2.3 PD Model for voxelotor and osivelotor

We aim to model how the effects of voxelotor and osivelotor on the percentage of *O*_2_ saturation change over time through PD analysis. The PD model for them involves a series of operations, as illustrated in Fig. 2C. Starting with the drug RBC concentrations obtained from the PK model, we sequentially transform these values into the percentage of hemoglobin occupancy and the percentage of hemoglobin modification, ultimately deriving the percentage of *O*_2_ saturation.

First, following the approach described in [48], we applied a unit conversion to transform the drug RBC concentration from *µg/ml* to *mM*. For voxelotor, the conversion was based on the linear relationship 337.5 *µg/ml* = 1 *mM*, as reported in [48], while for osivelotor, we used 386.4 *µg/ml* = 1 *mM*, according to [54].

Then, we performed a linear transformation to convert the drug RBC concentration ([*drug*]_*RBC*_) into the percentage of hemoglobin occupancy (% *Hb Occupancy*). For voxelotor, the transformation followed the equation from [48]:

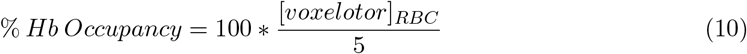

For osivelotor, the transformation was based on a linear relationship calibrated by fitting the PD data reported in [50]:

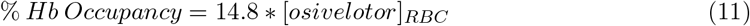

Next, we performed a linear transformation to obtain the corresponding percentage of hemoglobin modification(%*HbModification*) from the percentage of hemoglobin occupancy, using the following equation as adopted from [48].

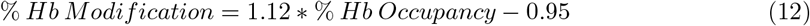

Finally, based on experimental data from [49], we interpolated linear relationships between the percentage of *O*_2_ saturation and the percentage of hemoglobin modification at liver and kidney/spleen oxygen level, respectively. These interpolated relationships were then used to predict *O*_2_ saturation across varying percentages of hemoglobin modification under liver and kidney/spleen oxygen levels.

#### 2.2.4 Kinetics Model

The employed kinetic model [32], grounded in classical nucleation theory, is designed to simulate the polymerization kinetics of HbS and predict the sickling of RBCs in SCD. As illustrated in Fig. 1, the model incorporates patient- and organ-specific factors—such as temperature, oxygen tension, mean corpuscular volume (MCV), mean corpuscular hemoglobin concentration (MCHC), and the mole fraction of various hemoglobin variants—to determine the number of nuclei formed through homogeneous and heterogeneous nucleation, as well as fiber growth and RBC sickling fraction as a function of time. To capture organ-specific sickling dynamics, we ran the RBC sickling kinetic model across a range of physiologically relevant oxygen partial pressures (*pO*_2_) corresponding to different tissues: liver (e.g., 30 mmHg) [55], spleen/kidney (e.g., 20 mmHg) [56, 57]. These values were used as fixed inputs in the polymerization kinetics equations, and the resulting fraction of sickled RBCs was computed accordingly for each condition. The simulation results estimate the reduction in the fraction of sickled RBCs. We first validated our PK/PD modeling framework and conducted simulations for single-drug scenarios involving either Bitopertin or voxelotor. We then extended our analysis to multi-drug scenarios, examining the combined use of HU and voxelotor, as well as Bitopertin and voxelotor, to explore whether simultaneous administration of multiple drugs in combination yields a greater therapeutic effect. Finally, we investigated the impact of noncompliance with Bitopertin or voxelotor, an issue commonly encountered in many SCD patients, to determine whether the consequences of noncompliance differ between the two drugs. We will use the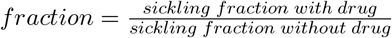 as a metric to evaluate the drug efficacy. Following the work of [25], we consider that a 70% decrease in the fraction of sickled RBCs under drug impact (relative sickling fraction*<*30%) is therapeutically effective.

## 3 Results

In this section, we present the outcomes of computational simulations based on integrated PK/PD and RBC sickling kinetic models. First, we calibrate PK/PD models to simulate the impact of Bitopertin, voxelotor and osivelotor using clinical data. Subsequently, we perform simulations to predict the efficacy of each drug in reducing the fraction of sickled RBCs across organ-specific oxygen levels, and different hematological conditions are evaluated. Then, we assess the therapeutic impact of combination therapies and evaluate the influence of treatment noncompliance on drug efficacy. The collective analysis of these results underscores the platform’s capacity for quantitatively assessing single and multi-agent anti-sickling therapies in a personalized and mechanistically informed manner.

### 3.1 Results of PK/PD modeling for Bitopertin

The results for the PK/PD modeling of Bitopertin are shown in Fig. 3. Fig. 3A shows the simulated Bitopertin plasma concentration over time for single dosages of 3 mg, 12 mg, and 24 mg. Fig. 3B displays the simulated Bitopertin plasma concentration for multiple dosages of 30 mg. Fig. 3C shows the simulated MCHC levels for multiple dosages of 10 mg, 30 mg, and 60 mg. The 50th percentile curves align well with the mean data, and the uncertainty bands adequately capture most of the observed data. The estimated parameters that result in these simulation outcomes are shown in Tab. 1.

**Table 1:** Estimated parameters of the PK and PD model for Bitopertin.

| Name | $V_1$ | $V_2$ | $K_a$ | $K_{12}$ | $K_{21}$ | $K_e$ | $E_{max}$ | $EC_{50}$ | $K_{out}$ |
| --- | --- | --- | --- | --- | --- | --- | --- | --- | --- |
| Unit | L | L | /hr | /hr | /hr | /hr | 1 | ng/ml | /hr |
| Value | 53.9 | 256 | 0.392 | 0.534 | 0.108 | 0.086 | 0.186 | 796 | $3.85e-4$ |

**Figure 3.**
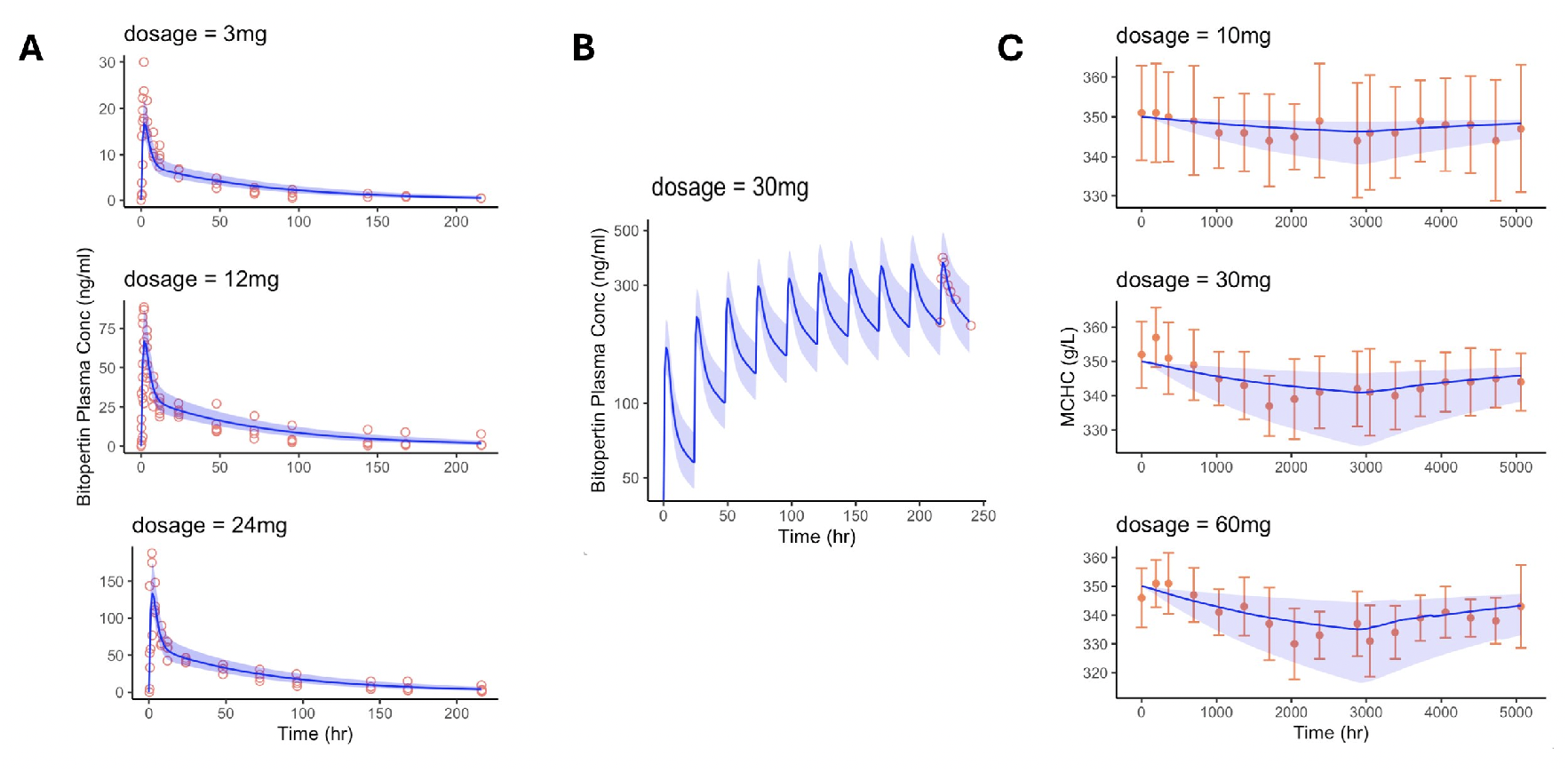
Calibration of the PK/PD modeling for Bitopertin. The blue solid lines represent the 50th percentile of the simulation results, while the blue ribbons indicate the 5th to 95th percentile range. (A) The simulated Bitopertin plasma concentrations over time for single dosages of 3 mg, 12 mg, and 24 mg. The red circles correspond to the observed data sourced from [45]. (B) The simulated Bitopertin plasma concentrations for a multiple-dose regimen of 30 mg administered once daily for 10 consecutive days. The red circles correspond to the observed data obtained from [46]. (C) The simulated mean corpuscular hemoglobin concentration (MCHC) levels for multiple dosages of 10 mg, 30 mg, and 60 mg. The red dots correspond to the mean values among the clinical data adopted from [47], and the error bars indicate the minimum and maximum values within the clinical data.

### 3.4 Impact of Bitopertin on reducing the fraction of sickled RBCs

Fig. 4 shows the effect of Bitopertin on reducing the fraction of sickled RBCs at organ-specific oxygen levels. At liver oxygen levels (Fig. 4A), the sickling fraction decreases gradually after Bitopertin administration, eventually reaching a plateau, with noticeable differences between various dosage levels. In contrast, at the kidney or spleen oxygen level (Fig. 4B), the impact of dosage is negligible. Figs. 4C and D further show that the trends of the relative sickling fraction of RBCs are consistent with results observed in Fig. 4A and B. These findings suggest that Bitopertin exhibits greater efficacy at higher oxygen levels, where baseline sickling is relatively low. Conversely, in hypoxic environments where the sickling fraction is near 1, increasing the dosage offers little additional benefit. Notably, only the 60 mg dosage under liver oxygen conditions achieves a reduction in sickling fraction approaching therapeutic effectiveness, implying that Bitopertin may have limited therapeutic effectiveness under physiological oxygenation conditions.

**Figure 4.**
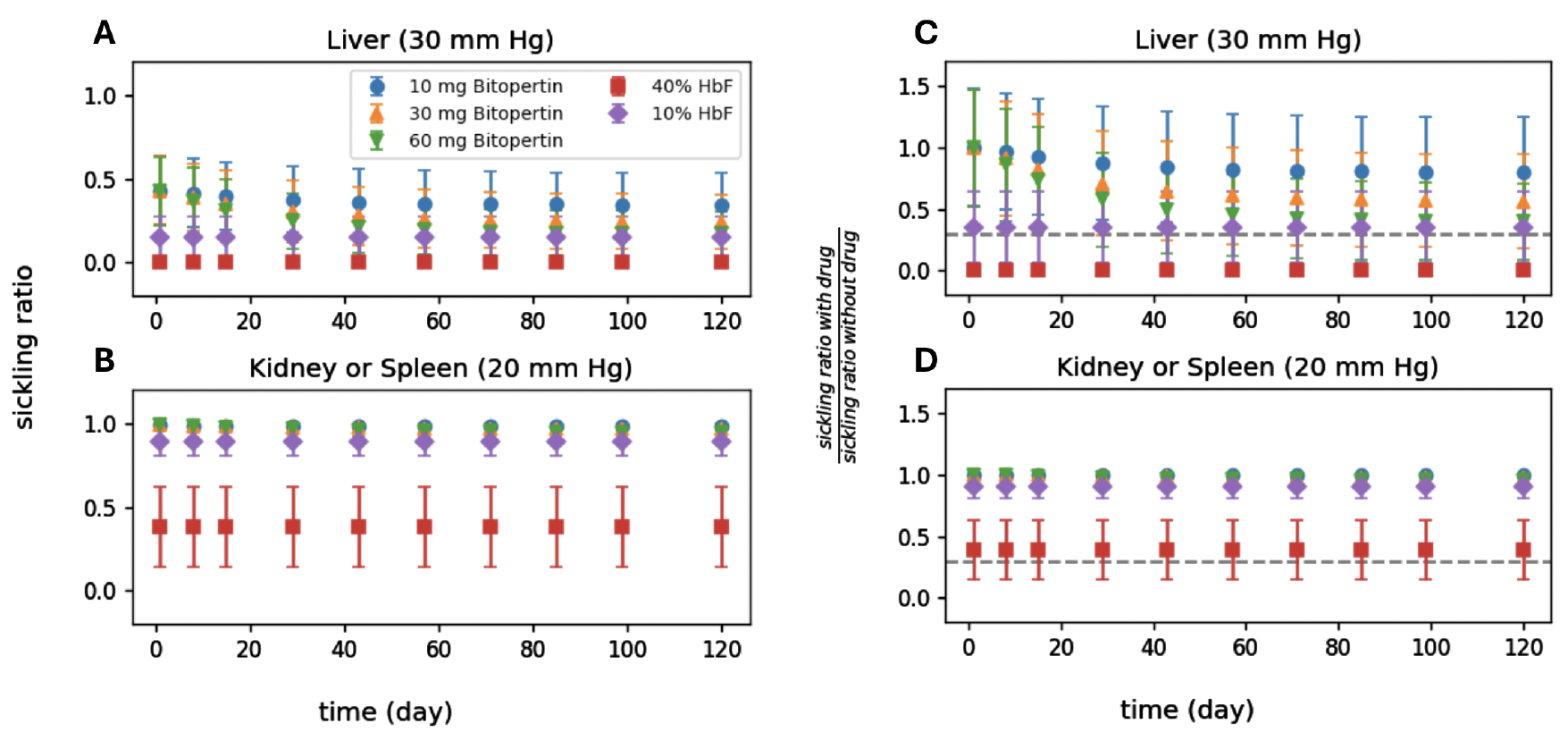
Impact of Bitopertin and HU on reducing the fraction of sickled RBCs at organ-specific oxygen levels. The impact of HU is incorporated by using 10% and 40% of HbF in the kinetic model, which are low and high ends of SCD patients’ responses to HU [58]. (A-B) show the absolute value of fractions of sickled RBCs, while (C-D) show the relative sickling fraction. The error bars result from 10 randomly sampled means and standard deviations of MCHC within the range of 32-36 g/dl and 0.7-2.5 g/dl, respectively. The grey dotted lines highlight a 70% decrease in the fraction of sickled RBCs under drug impact, which is considered therapeutically effective [25].

We further simulated the impact of Bitopertin on reducing the fraction of sickled RBCs at organ-specific oxygen levels across patients with different MCHC levels. Patients were categorized into three groups based on their MCHC levels: low (31–32 g/dL), mid (33–34 g/dL), and high (35–36 g/dL). At liver oxygen levels with low MCHC, predictions in Fig. 5A show that both the dosages of 30 mg and 60 mg could lead to near therapeutic efficacy. On the other hand, for patients with higher MCHC levels (Fig. 5C and E), none of the tested dosages achieve comparable therapeutic effects. Under more hypoxic conditions, Bitopertin fails to produce therapeutic efficacy across all MCHC groups, regardless of dosage. For comparison, we also simulate the sickling fraction of RBCs under HU, one of the FDA-approved drugs. The effect of HU was incorporated into the kinetic model by simulating HbF levels at 10% and 40%, representing the lower and upper response ranges observed in SCD patients receiving HU [58]. As shown in Fig. 5, when HbF levels induced by HU reach 40%, therapeutic efficacy is achieved across all MCHC groups under liver oxygen conditions. However, under kidney or spleen oxygen levels, the efficacy of HU diminishes with increasing MCHC and eventually falls below the therapeutic threshold.

**Figure 5.**
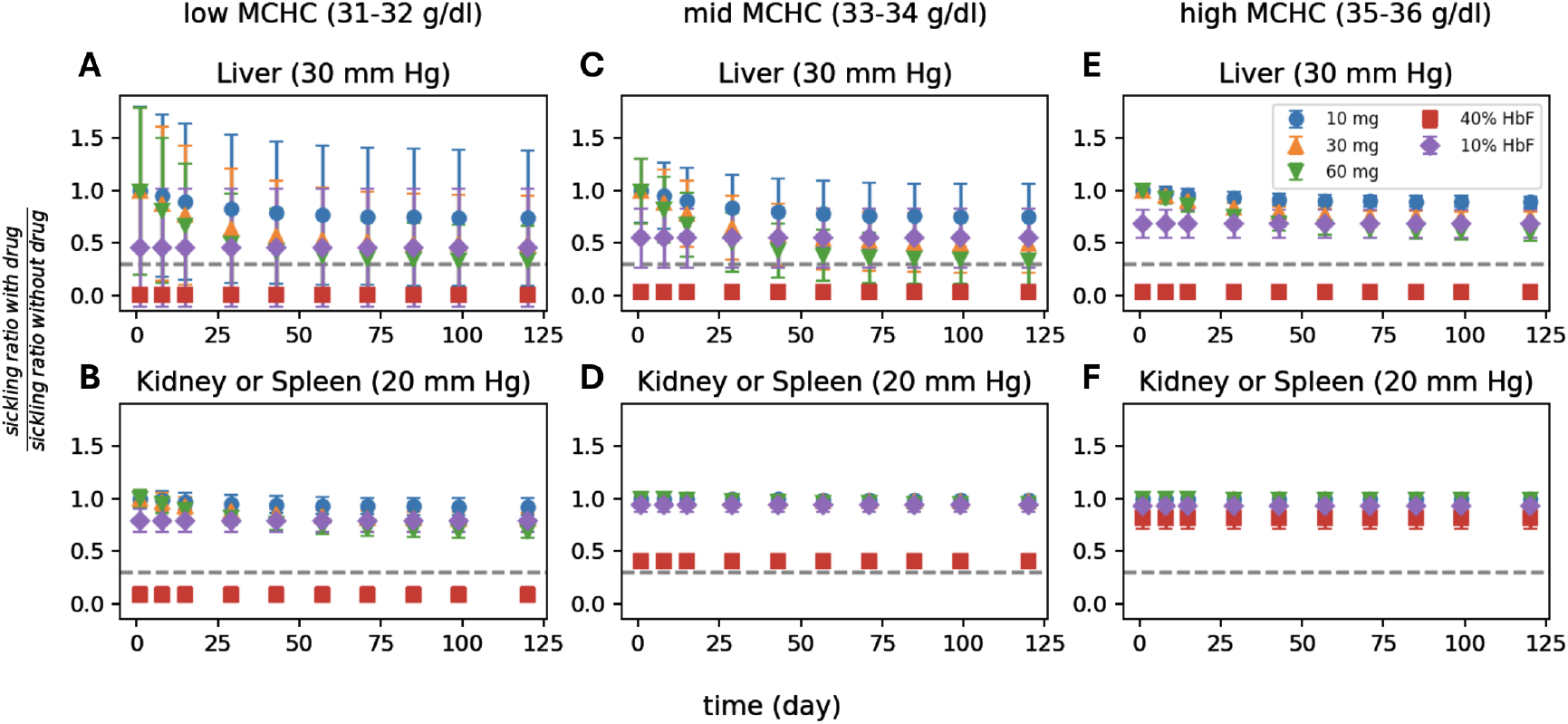
Impact of Bitopertin and HU on reducing the fraction of sickled RBCs at organ-specific oxygen levels for three MCHC groups, namely 31-32 g/dl (A-B), 33-34 g/dl (C-D), 35-36 g/dl (E-F). The impact of HU is incorporated by using 10% and 40% of HbF in the kinetic model, which are low and high ends of SCD patients’ responses to HU [58]. The error bars result from 10 randomly sampled means as specified in the figure and standard deviation within the range of 0.7-2.5 g/dl. The grey dotted lines highlight a 70% decrease in the fraction of sickled RBCs under drug impact, which is considered therapeutically effective [25].

These results suggest that patients with lower MCHC are more responsive to Bitopertin, particularly in organs with higher oxygen levels. In contrast, for patients with moderate to high MCHC or under hypoxic conditions, Bitopertin appears to be therapeutically ineffective. Comparatively, HU demonstrates broader and more consistent effectiveness, underscoring its clinical superiority over Bitopertin.

### 3.3 Results of PK/PD modeling for voxelotor and osivelotor

The results for the PK/PD modeling of voxelotor and osivelotor are shown in Fig. 6. Fig. 6A shows the PK simulations of voxelotor concentration within RBCs over time for single dosages of 100 mg, 1000 mg, 2000 mg, and 2800 mg, based on the observed data sourced from [48]. It also depicts the simulated mean osivelotor RBC concentrations over time for a single 100 mg dose, based on the observed data reported from [50]. Fig. 6B further displays the simulated voxelotor RBC concentration for multiple dosages of 300 mg, 600 mg, and 900 mg, calibrated using observed data sourced from [48]. The 50th percentile curves closely match the mean data, and the uncertainty bands effectively encompass the majority of the observed data. The estimated parameters that result in these simulation outcomes are summarized in Tab. 2.

**Table 2:**
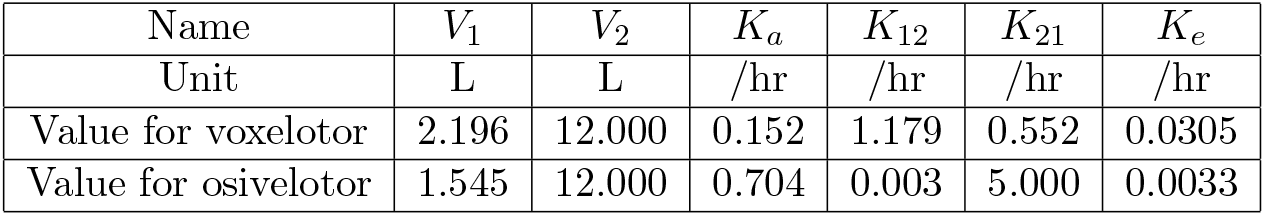
Estimated parameters of the PK model for voxelotor and osivelotor.

**Figure 6.**
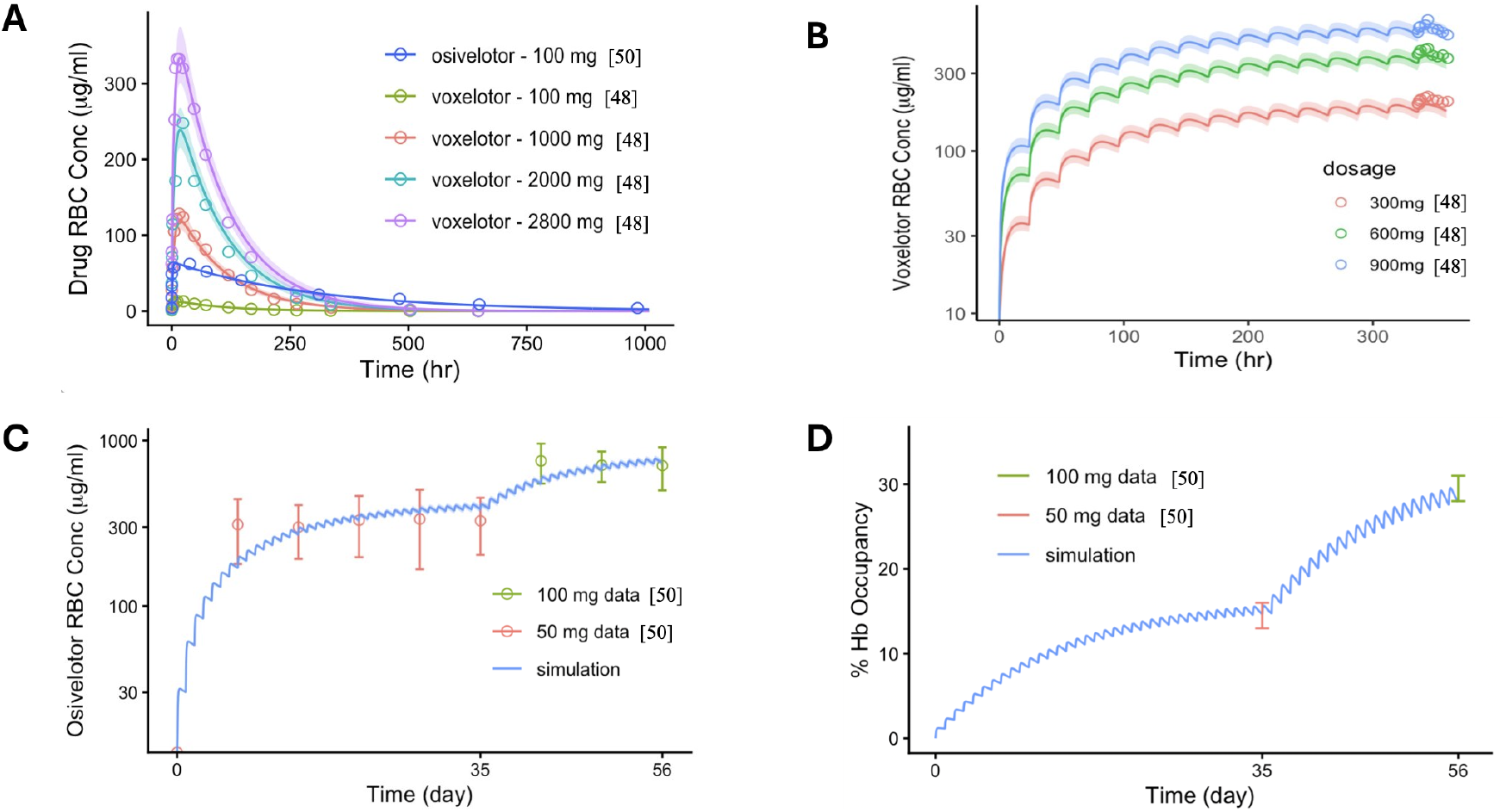
Calibration of the PK/PD modeling for voxelotor and osivelotor. (A) The simulated voxelotor RBC concentrations over time for single dosages of 100 mg, 1000 mg, 2000 mg, and 2800 mg, as well as the simulated osivelotor RBC concentrations over time for a single dosage of 100 mg. The solid lines represent the 50th percentile of the simulation results, while the ribbons indicate the 5th to 95th percentile range. The circles correspond to the observed data sourced from [48] for voxelotor and from [50] for osivelotor. (B) The simulated voxelotor RBC concentrations over time for multiple dosages of 300 mg, 600 mg, and 900 mg once every 24 hours for 15 days. The circles correspond to the observed data sourced from [48]. (C) The simulated mean osivelotor RBC concentrations over time for multiple dosages of 50 mg for five weeks, immediately followed by 100 mg for three weeks. The lines represent the simulation results, while the circles and error bars correspond to the clinical data reported from [50]. (D) The simulated mean percentage of hemoglobin occupancy following the same clinical treatment as in subfigure (C). The error bars represent the observed minimum and maximum ranges reported in [50].

Fig. 6C presents PK simulations of osivelotor concentrations within RBCs over time for a multiple dosing clinical treatment, based on the observed data sourced from [50]. The clinical treatment consisted of 50 mg administered daily for five weeks, immediately followed by 100 mg daily for three weeks. Fig. 6D displays the simulated mean percentage of hemoglobin occupancy following the same clinical treatment as in Fig. 6C. The simulations accurately capture the trend and align well with the clinical data.

### 3.4 Impact of voxelotor on reducing the fraction of sickled RBCs

Fig. 7 shows the effect of voxelotor in reducing the fraction of sickled RBCs at organ-specific oxygen levels. At liver oxygen levels (Fig. 7A), the sickling fraction decreases rapidly after voxelotor administration and quickly reaches a plateau, with no apparent difference observed between dosage levels. In contrast, at the kidney or spleen oxygen level (Fig. 7B), the impact of varying dosage is more pronounced. As shown in Figs. 7C and D, the trend of the relative sickling fraction is consistent with results shown in Fig. 7A and B. Figs. 7C and D further show that under liver oxygen conditions, all three examined dosages achieve therapeutic efficacy, whereas under kidney or spleen conditions, only the 1500 mg dosage reaches therapeutic levels. These results provide a mechanistic rationale for the selection of the 1500 mg dose in clinical trials for adult patients with SCD. Notably, the model’s prediction for the 900 mg dose shows efficacy comparable to *ex vivo* measurements at 20 mmHg reported for GBT021601 [59], a new agent that shares the same mechanism of action as voxelotor. These findings highlight the clinical relevance and predictive validity of our model.

**Figure 7.**
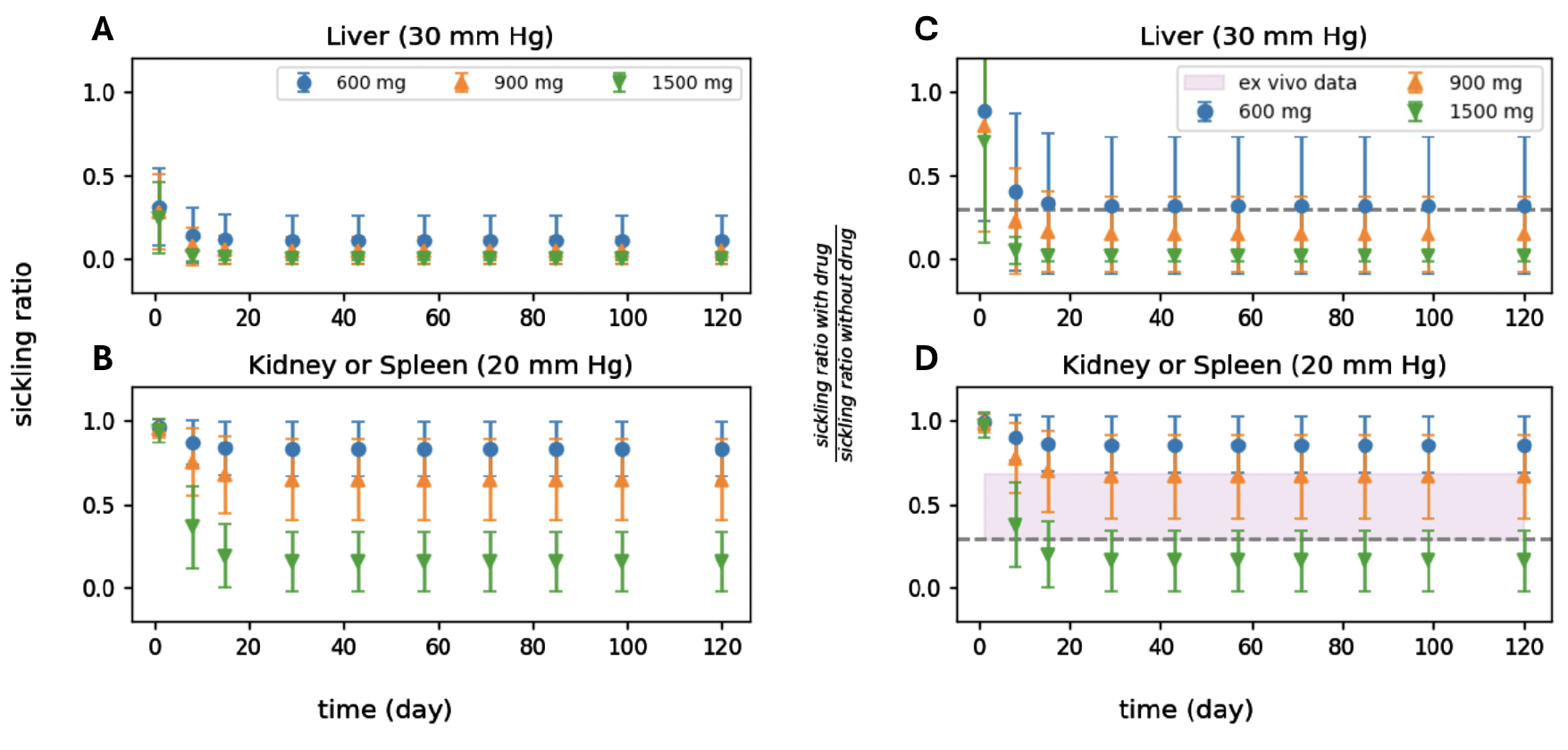
Impact of voxelotor on reducing the fraction of sickled RBCs at organ-specific oxygen levels. (A-B) show the absolute value of fractions of sickled RBCs, while (C-D) show the relative sickling fraction. The purple zone in (D) highlights the impact of GBT021601 measured *ex vivo* at 20 mmH [59]. GBT021601 is a new drug with the same mechanism of action as voxelotor. The error bars result from 10 randomly sampled means and standard deviations of MCHC within the range of 32-36 g/dl and 0.7-2.5 g/dl, respectively. The grey dotted lines highlight a 70% decrease in the fraction of sickled RBCs under drug impact, which is considered therapeutically effective [25]. 4 -12 years old, weighing 20 kg to 40 kg, will be provided with 900 mg daily during clinical trials. Weighing 10 kg to 20 kg will be provided with 600 mg daily.

We further simulated the effect of voxelotor on reducing the fraction of sickled RBCs at organ-specific oxygen levels across patients with different MCHC levels. As in the Bitopertin study, patients were divided into three groups based on their MCHC values: low (31–32 g/dL), mid (33–34 g/dL), and high (35–36 g/dL). Figs. 8A, C, and E show that under liver oxygen conditions, all three MCHC groups exhibit therapeutic efficacy, indicating that voxelotor is effective across a broad MCHC range in high-oxygen environments. Under kidney or spleen oxygen conditions (Figs. 8B, D, and F), therapeutic effectiveness is maintained for patients with low MCHC across all tested dosages. However, in the mid-MCHC and high-MCHC groups, only the 1500 mg dosage achieves therapeutic efficacy or near therapeutic efficacy. These results suggest that voxelotor’s efficacy is strongly influenced by organ-specific oxygen levels. Organs with relatively high oxygen levels are more responsive across all tested dosages, while those with low oxygen levels may require higher doses to achieve meaningful therapeutic outcomes. Consistently, the 1500 mg dose selected for clinical trials demonstrates efficacy across all MCHC groups for two different oxygen levels, supporting its rationale for use in adult SCD populations.

**Figure 8.**
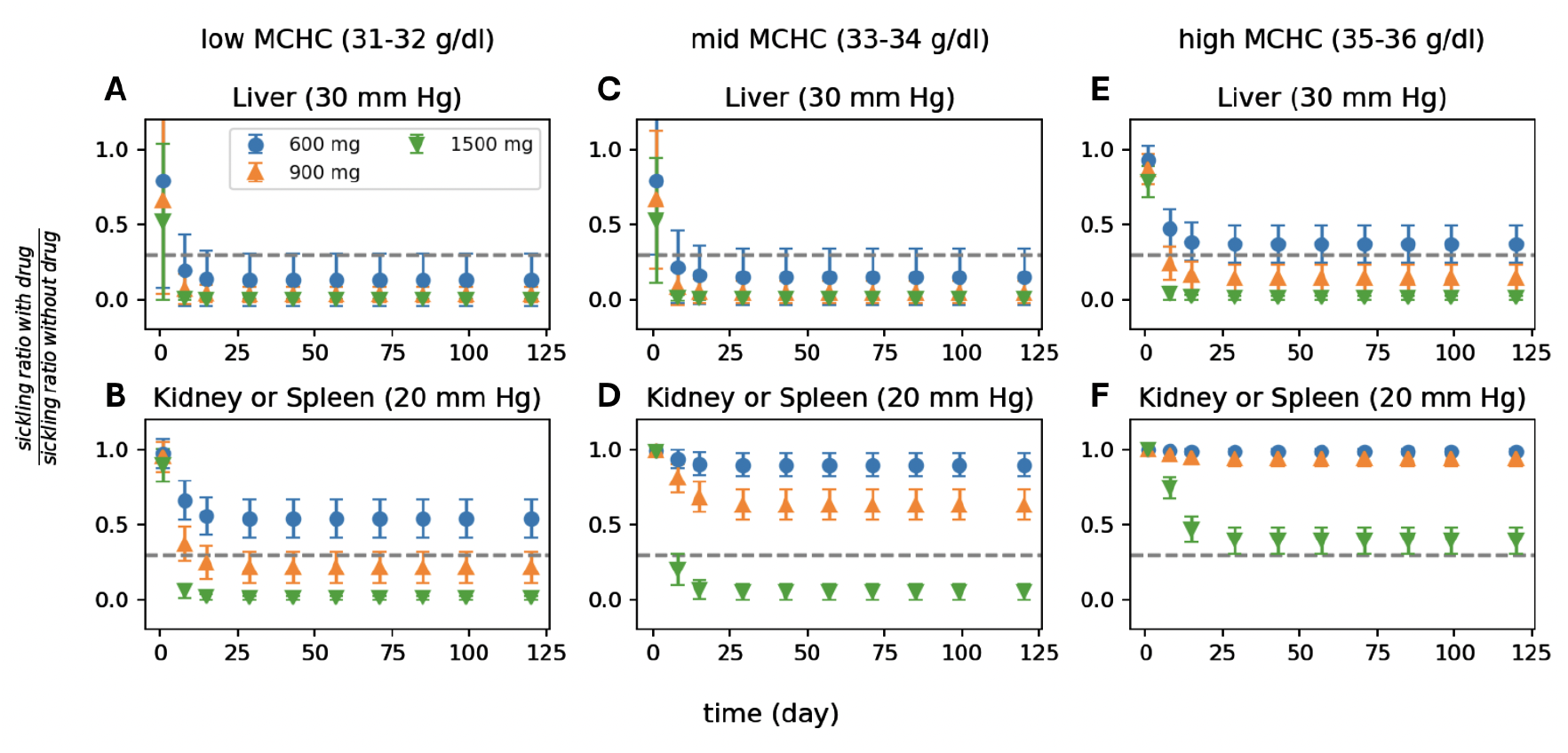
Impact of voxelotor on reducing the fraction of sickled RBCs at organ-specific oxygen levels for three MCHC groups, namely 31-32 g/dl (A-B), 33-34 g/dl (C-D), 35-36 g/dl (E-F). The error bars result from 10 randomly sampled means as specified in the figure and standard deviation within the range of 0.7-2.5 g/dl. The grey dotted lines highlight a 70% decrease in the fraction of sickled RBCs under drug impact, which is considered therapeutically effective [25].

### 3.5 Impact of multi-drugs on reducing the fraction of sickled RBCs

In this section, we simulated the impact of multi-drug therapy on reducing the fraction of sickled RBCs in two cases. In Case 1, we examined the combined effect of HU and voxelotor. Fig. 9 illustrates the impact of this combination on reducing the fraction of sickled RBCs at organ-specific oxygen levels. At liver oxygen levels (Fig. 9A, C, E, G), increasing HbF levels induced by HU have little to no effect on the efficacy of voxelotor across all three examined dosages. In contrast, at kidney or spleen oxygen level (Fig. 9B, D, F, H), the combined efficacy improves as HbF levels increase. These results suggest that the combination therapy of HU and voxelotor is likely to be more effective than voxelotor alone in organs with relatively low oxygenation. This observation supports findings from [60], where analysis of patient blood samples showed a significant reduction in sickled RBC fraction with increasing voxelotor dosage, even among patients already receiving HU therapy.

**Figure 9.**
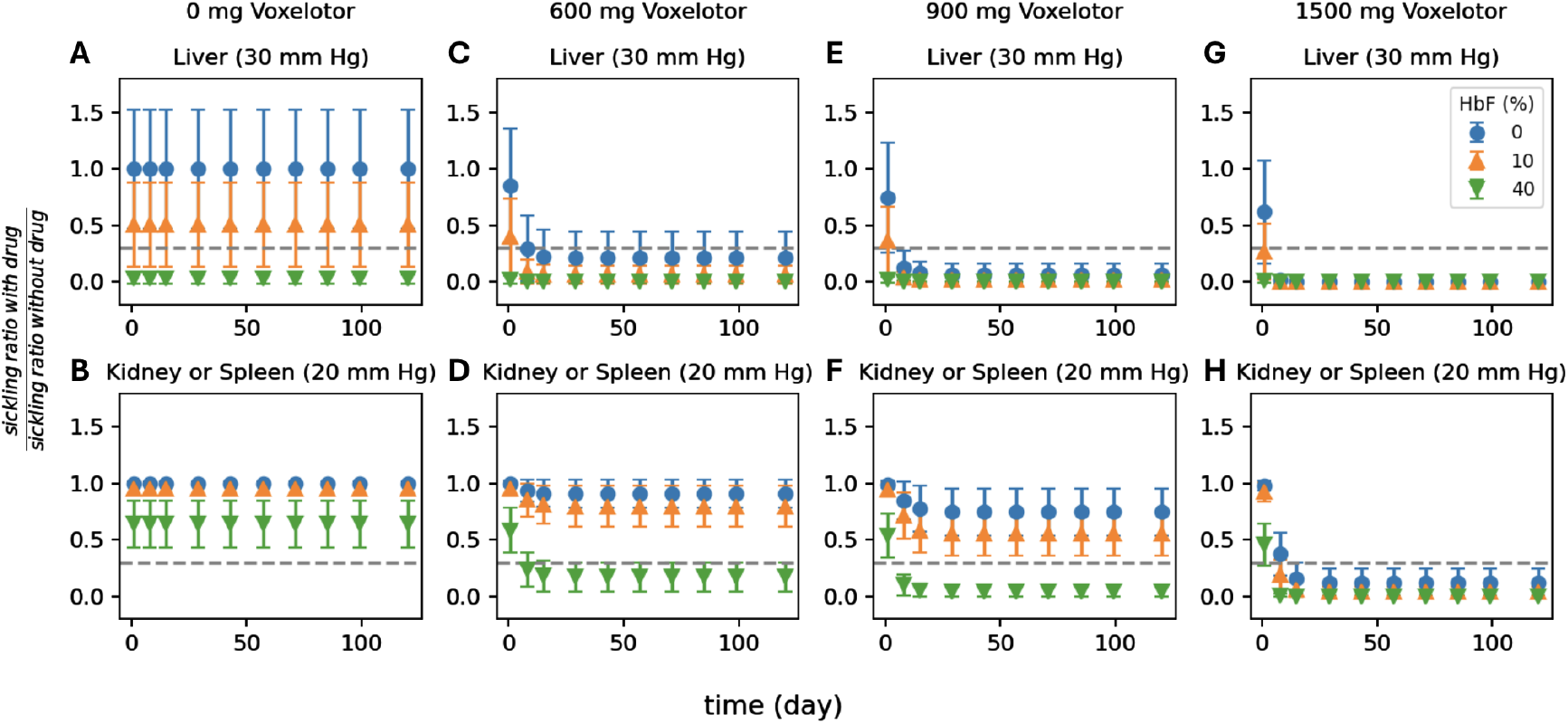
Impact of combined HU and voxelotor on reducing the fraction of sickled RBCs at organ-specific oxygen levels. The impact of HU is incorporated by using 10% and 40% of HbF in the kinetic model, which are low and high ends of SCD patients’ responses to HU [58]. The error bars result from 10 randomly sampled means and standard deviations of MCHC within the range of 32-36 g/dl and 0.7-2.5 g/dl, respectively. The grey dotted lines highlight a 70% decrease in the fraction of sickled RBCs under drug impact, which is considered therapeutically effective [25].

In Case 2, we investigated the combined effect of Bitopertin and voxelotor on reducing the fraction of sickled RBCs at organ-specific oxygen levels. As shown in Fig. 10A, in the absence of voxelotor, treatment with Bitopertin leads to a gradual decrease in sickling, eventually reaching a plateau. The difference in drug efficacy between various Bitopertin dosages is noticeable. However, when the voxelotor dosage increases to 600 mg, 900 mg, and 1500 mg (Fig. 10C, E, G), the drug efficacy follows a similar trend but declines much more rapidly after administration, ultimately reaching a plateau near zero. These observations indicate that the combined drug efficacy is primarily driven by voxelotor. A similar pattern is observed at the kidney or spleen oxygen level (Fig. 10B, D, F, H), where the influence of voxelotor on drug efficacy overwhelms that of Bitopertin. Overall, when Bitopertin and voxelotor are combined, the modeled drug efficacy is largely dominated by voxelotor, with only minimal additional benefit from increasing the Bitopertin dosage.

**Figure 10.**
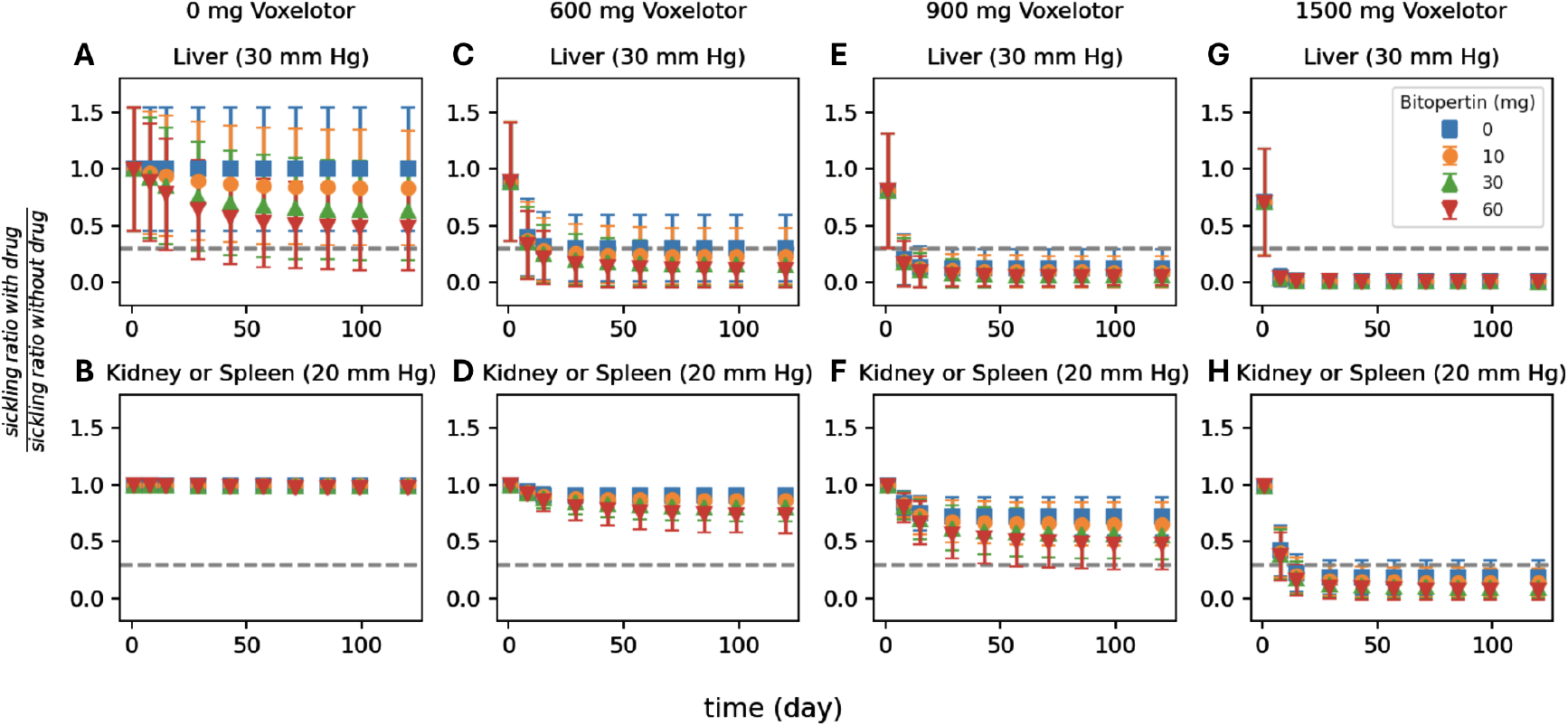
Impact of combined Bitopertin and voxelotor on reducing the fraction of sickled RBCs at organ-specific oxygen levels. The error bars result from 10 randomly sampled means and standard deviations of MCHC within the range of 32-36 g/dl and 0.7-2.5 g/dl, respectively. The grey dotted lines highlight a 70% decrease in the fraction of sickled RBCs under drug impact, which is considered therapeutically effective [25].

### 3.6 Impact of drug noncompliance on altering the fraction of sickled RBCs

Drug compliance with prescribed treatment is critically important in the management of SCD. Poor compliance can significantly compromise treatment effectiveness, increase the risk of developing vaso-occlusion and other complications [61–63]. In this section, we employed our framework to investigate the impact of drug noncompliance on the dynamics of the sickled RBC fraction.

Figs. 11A and B present simulation results for missed Bitopertin doses (between days 21–25) after the drug’s anti-sickling effect has reached steady state under liver and kidney oxygen levels. Notably, even after a five-day noncompliance, the increase in the fraction of sickled RBCs is insignificant despite the fact that Bitopertin is not therapeutically effective. The minimal short-term impact of noncompliance is likely attributable to the mechanism by which Bitopertin modulates MCHC and MCV of RBCs, primarily through its influence on erythropoiesis, a process that evolves gradually over time and the drug effects also decay relatively slowly. On the other hand, Fig. 11C shows that noncompliance to voxelotor produces a small increase in the fraction of sickled RBCs in organs with relatively high oxygen levels. In contrast, Fig. 11D shows that missing just two days of voxelotor treatment leads to a rapid elevation of the portion of sickled RBCs under more hypoxic conditions, such as those found in the kidney or spleen. These findings provide a mechanistic explanation for clinical observations of “rebound” hemolysis and organ dysfunction associated with voxelotor noncompliance [64].

**Figure 11.**
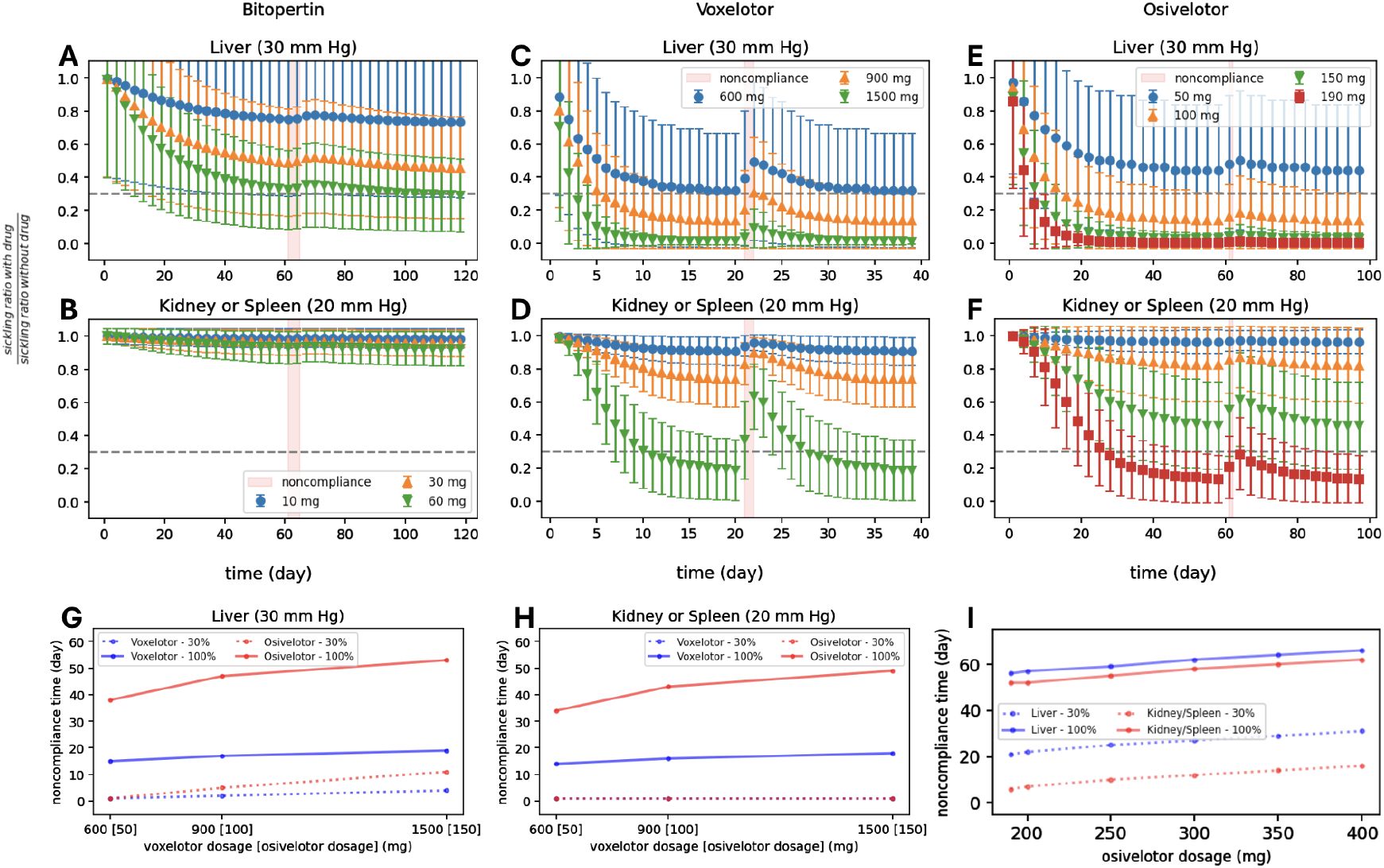
Impact of noncompliance on altering the dynamics of the impact of Bitopertin, voxelotor, and osivelotor. In the case of Bitopertin (A-B), noncompliance occurs from day 61 to 65, and daily medication is resumed thereafter. In the case of voxelotor (C-D), the noncompliance occurs from day 21-22, and daily medication is resumed thereafter. In the case of osivelotor (E-F), the noncompliance occurs from day 81-82, and daily medication is resumed thereafter. The error bars result from 10 randomly sampled means and standard deviations of MCHC within the range of 32-36 g/dl and 0.7-2.5 g/dl, respectively. The light red zone highlights the noncompliance period. The grey dotted lines highlight a 70% decrease in the fraction of sickled RBCs under drug impact, which is considered therapeutically effective [25]. (G-H) Noncompliance duration required for voxelotor and osivelotor to reach scaled sickling ratios exceeding 30% and 100% across clinical trial dosage levels. (I) Noncompliance duration required for osivelotor to reach scaled sickling ratios exceeding 30% and 100% across extended dosage levels.

We further examined the impact of noncompliance on osivelotor and compared it with voxelotor. First, three dosage levels, namely 50 mg, 100 mg, and 150 mg, were simulated following the clinical trial design in [50]. As shown in Figs. 11E and F, after the fraction of sickled RBCs reached a steady state at approximately 60 days, missing two consecutive doses of osivelotor led to smaller increases in the fraction of sickled RBCs compared with those of voxelotor. This finding indicates that osivelotor exhibits greater tolerance to short-term noncompliance under both liver and kidney/spleen oxygen conditions. However, we noted that at these three examined dosages, osivelotor only achieves therapeutic efficacy under liver oxygen levels, not under the lower oxygen levels representative of kidney and spleen environments. Therefore, we further increased the simulated dosage to 190 mg. As shown by the red curves in Figs. 11E and F, at this dosage, the fraction of sickled RBCs decreased below the therapeutic threshold, and missing two consecutive doses did not cause the fraction to rise above the 30% therapeutic limit. These results highlight the critical importance of compliance with voxelotor and osivelotor therapy for maintaining clinical efficacy.

Next, we examine the relationship between dosage and noncompliance-driven increases in sickled RBC percentage. The association between drug dosage and the duration of noncompliance that leads to loss of therapeutic efficacy for voxelotor and osivelotor, was summarized in Figs. 11G and H. For comparison, we used therapeutic dosages following the clinical trials [48, 50]: 600 mg, 900 mg, and 1500 mg for voxelotor, and 50 mg, 100 mg, and 150 mg for osivelotor. Across all evaluated dosage pairs, osivelotor exhibited substantially longer times to reach scaled sickling ratios exceeding 30% (failing to reach therapeutic anti-sickling effect) and 100% (zero anti-sickling effect), indicating markedly greater resilience to noncompliance than voxelotor. We further examined the noncompliance tolerance of osivelotor across extended dosage levels (190–400 mg). As shown in Fig. 11I, osivelotor displays a tolerance of approximately one week or much longer at these higher dosages under both liver and kidney/spleen oxygen levels. These results could provide guidance for optimizing dosage strategies to mitigate the risks associated with noncompliance.

## 4 Discussion and Summary

In this study, we present a computational platform that combines PK/PD models with a kinetic model of RBC sickling in critical tissues, enabling efficient prediction of dosage-dependent therapeutic efficacy for various anti-sickling agents. This platform enables patient-specific predictions based on hematological factors and organ-specific oxygen levels. To validate its effectiveness, we first applied our platform to assess the therapeutic efficacy of two FDA-approved drugs, HU and voxelotor. We can clearly observe the impact of the two drugs on reducing the sickling fraction of the RBCs under different drug doses and organ-specific oxygen levels. Notably, the drug efficacy varies significantly in patients with different levels of MCHC, underscoring the importance of delivering patient-specific dosing to achieve optimal therapeutic outcomes. Overall, our results show that both HU and voxelotor present clear anti-sickling effects in most tested scenarios, supporting the outcomes reported in their clinical trials [22, 65, 66]. It is noted that in our simulations, we focus solely on the effect of HU in preventing RBC sickling. However, previous studies [67–70] have shown that HU can also ameliorate the symptoms of SCD through multiple additional mechanisms, including reducing RBC adhesion and attenuating inflammatory responses.

We then extended our analysis to two potential anti-sickling candidates, namely Bitopertin and osivelotor. Bitopertin, which is currently under clinical investigation for erythropoietic protoporphyria [71, 72], has been reported to reduce RBC MCHC by modulating the erythropoiesis process, thereby indirectly mitigating sickling. Our simulations reveal that while Bitopertin exerts a modest protective effect against RBC sickling, its overall efficacy remains substantially lower than that of HU and voxelotor, suggesting limited therapeutic potential for direct treatment of SCD. In contrast, our model predicts that osivelotor, the next-generation analog of voxelotor with improved PK properties [50, 54], can achieve comparable anti-sickling efficacy as voxelotor at markedly lower doses. This finding highlights osivelotor’s strong potential as new treatment option for SCD. Collectively, these results underscore the translational capability of our computational platform as an *in silico* tool for post-screening of emerging anti-sickling therapies.

Additionally, we demonstrated the versatility of our platform by predicting the anti-sickling effects of multi-drug therapies. Specifically, we examined two combination regimens: HU with voxelotor, and Bitopertin with voxelotor. Our results indicate that the HU–voxelotor combination is likely more effective than voxelotor alone in organs with relatively low oxygenation. This finding supports clinical observations reported in [60], which showed a significant reduction in sickled RBC fractions with increasing voxelotor doses among patients already receiving HU therapy. In contrast, when Bitopertin and voxelotor are combined, our model predicts that the therapeutic effect is largely driven by voxelotor, with only minimal added benefit from increasing Bitopertin dosage. These results highlight the model’s capability to evaluate the combined efficacy of multi-agent therapies, which are often challenging to assess directly through clinical trials due to complexity, cost, and patient recruitment constraints.

We also employed our model to investigate the consequences of noncompliance with Bitopertin, voxelotor, and osivelotor. Our results indicate that while the impact of noncompliance is minimal for Bitopertin, it can substantially compromise the therapeutic efficacy of both voxelotor and osivelotor. Specifically, the model predicts that even a two-day interruption of voxelotor therapy can cause a rapid rise in the fraction of sickled RBCs to levels insufficient to maintain therapeutic benefit in organs such as the kidney and spleen, where oxygen tension is relatively low. This mechanistic insight provides an explanation for the clinically observed “rebound” hemolysis and organ dysfunction associated with voxelotor noncompliance [64]. Moreover, our findings may help interpret clinical reports of increased mortality and VOC rates among voxelotor-treated patients that ultimately contributed to its market withdrawal. As illustrated in Fig. 11D, the abrupt increase in RBC sickling upon treatment interruption may be more deleterious than in untreated conditions, as the circulatory system lacks sufficient time to adapt to the sudden surge in sickled cells, resulting in a higher incidence of VOC compared to patients receiving a placebo. Our simulation results further demonstrated that osivelotor could enhance the tolerance to noncompliance, reducing the risk of experiencing sickling-induced adverse events. We further quantified the relationship between drug dosage and the duration of noncompliance that leads to loss of therapeutic efficacy for voxelotor and osivelotor. This analysis could provide guidance for optimizing dosage strategies to mitigate the risks associated with treatment interruptions and ensure sustained therapeutic efficacy.

Despite the strengths of our framework, there are a few limitations to acknowledge. First, we assumed all the RBCs are deoxygenated to a fixed organ-specific oxygen tension, whereas *in vivo* oxygen levels may fluctuate due to perfusion dynamics and patient-specific physiological changes.

In addition, the oxygen levels of individual RBCs within each organ may not be the same. Second, although MCHC variability was considered, other important factors such as patient age, disease severity, residual spleen function [73], and comorbidities were not explicitly incorporated into the model. Third, for combination therapy simulations, we did not have experimentally validated data to benchmark our predictions. Moreover, we assumed additive drug effects and did not account for potential PK or PD interactions between co-administered agents, which may influence efficacy or toxicity in real-world settings. For example, a prior clinical trial suggested that HU and voxelotor were not synergistic in terms of clinical outcomes of the HOPE study [74], and the reason is still not well-understood. Recent AI methods in PK modeling—such as AI-Aristotle [75] and CMINNs [76] for compartment models—specifically reveal that PK and PD model parameters can be captured by Physics-Informed Networks [77, 78] if they are treated as time-varying. This approach has been shown to enhance the accuracy of model fitting and improve the prediction of dose-response effects. Although we have considered these parameters constant in this work, pursuing this direction in future work may enable even more accurate and robust predictive modeling.

In summary, our *in silico* platform serves as a valuable tool for post-screening analysis of potential anti-sickling agents by considering their PK and anti-sickling efficacy under patient-specific hemoglobin level and organ-specific oxygen level, thereby gaining insights into their potential therapeutic efficacy alone or in combination before clinical trials. It is particularly well-suited for analyzing repurposed drugs, leveraging existing PK data from previous clinical trials for other conditions. This comprehensive modeling approach can be further extended to evaluate other anti-sickling agents and optimize dosing strategies to improve clinical outcomes. Future work may focus on refining model assumptions, incorporating patient variability, and expanding the model to capture additional biological mechanisms related to SCD.

## Acknowledgments

H.L. would like to thank Will Savage, Akshay Buch, Paul Hinderliter, Marcus Carden, and John Chapin from DiscMedicine for reviewing the manuscript.

